# Proinflammatory cytokine-induced alpha-cell impairment in human islet microtissues is partially restored by dual incretin receptor agonism

**DOI:** 10.1101/2024.06.26.600877

**Authors:** Kristine Henriksen, Chantal Rufer, Alexandra C. Title, Sayro Jawurek, Bolette Hartmann, Jens J. Holst, Filip K. Knop, Burcak Yesildag, Joachim Størling

**Affiliations:** Translational Type 1 Diabetes Research, Department of Clinical and Translational Research, Steno Diabetes Center Copenhagen, Herlev, Denmark; InSphero AG, Schlieren 8952, Switzerland; Department of Biomedical Sciences, Faculty of Health and Medical Sciences, University of Copenhagen, Copenhagen, Denmark; Novo Nordisk Foundation Center for Basic Metabolic Research, Faculty of Health and Medical Sciences, University of Copenhagen, Copenhagen, Denmark; Center for Clinical Metabolic Research, Gentofte Hospital, University of Copenhagen, Hellerup, Denmark; Department of Clinical Medicine, Faculty of Health and Medical Sciences, University of Copenhagen, Copenhagen, Denmark; Translational Type 2 Diabetes Research, Department of Clinical and Translational Research, Steno Diabetes Center Copenhagen, Herlev, Denmark

**Keywords:** GIP, GLP-1, Glucagon secretion, Human model, Incretins, Liraglutide, Pancreatic alpha cells, Proinflammatory cytokines, Type 1 diabetes

## Abstract

**Aims/hypothesis:** In type 1 diabetes, the counterregulatory glucagon response to low plasma glucose is impaired. The resulting increased risk of hypoglycaemia necessitates novel strategies to ameliorate alpha-cell impairment. Here, we aimed to establish an in vitro model of alpha-cell impairment in type 1 diabetes using human islet microtissues (MTs) exposed to proinflammatory cytokines. Additionally, we investigated the therapeutic potential of incretin receptor agonists in improving alpha-cell responses to low glucose.

**Methods:** Human islet MTs were exposed to proinflammatory cytokines (IL-1β, IFN-γ, and TNF-α) for 1 day (short-term) and 6 days (long-term). Alpha-cell function was assessed by sequential glucose-dependent secretion assays at 2.8 and 16.7 mmol/l glucose, followed by glucagon measurements. Additional evaluations included ATP content, caspase-3/7 activity, chemokine secretion, and expression of islet transcription factors and hormones. The effects of incretin receptor agonist treatment (glucose-dependent insulinotropic polypeptide (GIP) analogue [D-Ala^2^]-GIP ± liraglutide) alongside or after cytokine exposure were also investigated, focusing on low glucose-dependent glucagon secretion.

**Results:** Short-term cytokine exposure increased glucagon secretion at both 2.8 and 16.7 mmol/l glucose. In contrast, long-term cytokine exposure caused dose-dependent suppression of glucagon secretion at 2.8 mmol/l glucose, resembling a type 1 diabetes phenotype. Long-term cytokine exposure also diminished somatostatin secretion, reduced ATP content, increased caspase 3/7 activity, and decreased islet transcription factor and hormone expression. Despite cytokine-induced impairment, alpha cells partially retained secretory capacity to L-arginine stimulation. Treatment with incretin receptor agonists during long-term cytokine exposure did not prevent alpha-cell impairment. However, acute treatment with [D-Ala^2^]-GIP ± liraglutide or the single-molecule dual agonist tirzepatide partially restored glucagon secretion at low glucose.

**Conclusions/interpretation:** Long-term cytokine exposure of human islet MTs impaired glucagon secretion to low glucose, creating a type 1 diabetes alpha-cell phenotype. This cytokine-induced alpha-cell impairment was partially restored by [D-Ala^2^]-GIP ± liraglutide and tirzepatide, respectively.

**Research in context:** *What is already known about this subject?:*

- The counterregulatory alpha-cell response to low glucose is impaired in type 1 diabetes, increasing the risk of hypoglycaemia.
- Limited translatability of rodent islet findings highlights the need for human islet models.
- Actions of the incretin hormones glucose-dependent insulinotropic polypeptide (GIP) and glucagon-like peptide 1 (GLP-1) have mainly been studied in the context of type 2 diabetes and hyperglycaemia but less in type 1 diabetes and hypoglycaemia.

*What is the key question?:*

- Can alpha-cell impairment in type 1 diabetes be modelled in vitro by exposing human islet microtissues (MTs) to proinflammatory cytokines, and could incretins protect against this?

*What are the new findings?:*

- Long-term (6-day) exposure to proinflammatory cytokines produces a type 1 diabetes phenotype of alpha-cell impairment to low glucose in islet MTs
- Acute dual treatment with incretin receptor agonists partially restored glucose-dependent glucagon secretion in cytokine-exposed islet MTs — an effect mainly carried by GIP receptor agonism and not opposed by GLP-1 receptor agonism.

*How might this impact on clinical practice in the foreseeable future?:*

- Investigating incretin receptor agonists in a preclinical in vitro model of alpha-cell impairment may reveal their potential and fast-track their use as safeguards against hypoglycaemia in type 1 diabetes.

## Introduction

The alpha cells of the pancreatic islets of Langerhans play a pivotal role in glucose homeostasis by secreting counterregulatory glucagon in response to low glucose levels, which stimulates hepatic glucose output to prevent hypoglycaemia [1, 2]. In individuals with type 1 diabetes, this glucose-dependent counterregulatory alpha-cell response is impaired, while their responses to arginine persist. Consequently, individuals with type 1 diabetes are at an increased risk of hypoglycaemia [3–6]. Despite significant advances in insulin therapy, hypoglycaemia remains a critical challenge in type 1 diabetes management by limiting proper glycaemic control and by contributing to morbidity and mortality [7, 8]. Moreover, the risk of hypoglycaemia negatively impacts the quality of life of individuals with type 1 diabetes and their relatives [9]. Still, the mechanisms underlying glucose-dependent regulation of glucagon secretion in health and type 1 diabetes pathology remain incompletely understood [10, 11]. Efforts to elucidate alpha-cell impairment in the context of type 1 diabetes are hampered by the limited availability of human islets from individuals with type 1 diabetes [12, 13].

Exposure of isolated islets to diabetogenic proinflammatory cytokines is a widely used model of immune-mediated beta-cell destruction [14–17]. However, whether cytokine-exposed islets are also a valid model of human alpha-cell impairment in response to low glucose has yet to be established. Of note, human islets exhibit considerable heterogeneity, presenting challenges for studies of alpha-cell responses in vitro [18–22]. Recent advances have led to the development of reaggregated islet models, also termed islet microtissues (MTs), offering improved uniformity in cellular composition and size [23–26]. Reaggregated human islets thus provide a valuable platform for investigating islet function in a standardised setting, enhancing the ability to draw biologically meaningful conclusions while maintaining the physiological relevance inherent to human donors as opposed to rodent models and cell lines [23–26]. This is particularly relevant in the context of preclinical screening and validation of novel therapeutic interventions [27].

The gut hormones glucagon-like peptide-1 (GLP-1) and glucose-dependent insulinotropic polypeptide (GIP) play essential roles in glucose homeostasis through their glucose-dependent effects on islet insulin and glucagon secretion and have been proposed for treatments to enhance overall glycaemic control in individuals with type 1 diabetes [28, 29]. GLP-1 receptor agonists (GLP-1RAs) are extensively used in the treatment of type 2 diabetes and are also emerging as a promising therapy to maintain residual beta-cell function in newly diagnosed type 1 diabetes [30]. The glucagonotropic effect of GIP at low glucose concentrations holds promise to reduce the risk of hypoglycaemia in type 1 diabetes [31], though GIP alone may not be sufficiently effective [32]. Single-molecule dual GIP and GLP-1 receptor agonists are currently under extensive development and are considered crucial for the advancement of diabetes management, with tirzepatide already approved to date [33, 34], though their effects on alpha-cell function and hypoglycaemia are often overlooked. Furthermore, species-specific differences in receptor engagement underscore the necessity of using human models to accurately evaluate their therapeutic efficacy and potential in type 1 diabetes and to elucidate signalling pathways for the identification of more specific therapeutic targets [35, 36].

We here present an in vitro model of alpha-cell dysfunction in type 1 diabetes involving human islet MTs exposed to proinflammatory cytokines. Additionally, as a proof-of-concept, we investigate the effects of single and dual incretin receptor agonism using [D-Ala^2^]-GIP (a GIPRA) and liraglutide (a GLP-1RA) to evaluate their therapeutic potential for improving the glucagon response to low glucose in type 1 diabetes.

## Methods

### Human islet microtissues

Human islet microtissues (MTs), also known as reaggregated islets, were acquired from InSphero AG (Zurich, Switzerland) [23, 24]. Islet MTs from a total of 12 donors (M/F 11/1, age 49 ± 11 years, BMI 29 ± 3.6 kg/m^2^, HbA1c 5.3 ± 0.4%) formed the basis of this study. All islet MTs were sourced from non-diabetic, anonymised, and cadaveric organ donors. Informed consent was obtained at the isolation site according to local ethical legislation. Donor details are summarised in Table 1. Islet MTs were maintained in 3D InSight Human Islet Maintenance Medium (InSphero AG, Switzerland) at 37°C in a humidified 5% CO2 incubator, with the culture medium renewed every 2–3 days.

**Table 1.** Human islet donor details. Based on the ‘checklist for reporting human islet preparations used in research’ adapted from [69].

| Donor | Figures | Unique identifier | Age (years) | Sex (M/F) | BMI (kg/m<sup>2</sup>) | HbA1c (%) | Source of islet MTs<sup>a</sup> | Islet isolation center<sup>b</sup> | History of diabetes |
| --- | --- | --- | --- | --- | --- | --- | --- | --- | --- |
| 1 | Fig. 1b<br>ESM Fig. 1a, 2d | AKAR092 | 55 | M | 30.4 | 5.6 | InSphero | Prodo Laboratories | No |
| 2 | Fig.1c<br>ESM Fig. 1b, 2e | AKBL227 | 60 | M | 32.9 | 5.0 | InSphero | Prodo Laboratories | No |
| 3 | Fig. 2b, 3a<br>ESM Fig. 2a | AJG3470 | 46 | M | 23.1 | 4.9 | InSphero | Prodo Laboratories | No |
| 4 | Fig. 2c-d, 3b<br>ESM Fig. 2b, 3a-j | AJHU080 | 51 | M | 31.3 | 5.5 | InSphero | Prodo Laboratories | No |
| 5 | Fig. 2e-f, 3c<br>ESM Fig. 2c | AJIS384 | 67 | M | 34.9 | 5.3 | InSphero | Prodo Laboratories | No |
| 6 | ESM Fig. 2b, 2f | AJHR202 | 54 | M | 28.2 | 5.7 | InSphero | Prodo Laboratories | No |
| 7 | ESM Fig. 2g<br>Fig. 7e-f, 8a-b | AKCE427 | 48 | M | 23.7 | 5.5 | InSphero | Prodo Laboratories | No |
| 8 | Fig. 3d-e, 4, 5a-c, 7b<br>ESM Fig. 4 | AJI4346 | 37 | M | 29.1 | 5.4 | InSphero | Prodo Laboratories | No |
| 9 | Fig. 3e, 3g-i | AKJG366 | 58 | F | 31.0 | 4.7 | InSphero | Prodo Laboratories | No |
| 10 | Fig. 3f, 3j-m | AKJ1361 | 47 | M | 28.1 | 5.3 | InSphero | Prodo Laboratories | No |
| 11 | Fig. 5e-h, 6a-j, 7c-d<br>ESM Fig. 5a-j | AJKO093 | 32 | M | 26.9 | 4.9 | InSphero | Prodo Laboratories | No |
| 12 | Fig. 8c-d | AKHX448 | 33 | M | 33.1 | 5.7 | InSphero | Prodo Laboratories | No |

<sup>a</sup>InSphero AG, Switzerland. <sup>b</sup>Prodo Laboratores Inc, USA.

### Proinflammatory cytokine treatment

Islet MTs were cultured for 1 day (short-term) or 6 days (long-term) with or without a cytokine mix of recombinant human IL-1β (R&D Systems, USA or Sigma-Alrich, USA), recombinant human IFN-γ (PeproTech, USA or R&D Systems, USA) and recombinant human TNF-α (R&D Systems, USA) in a 1:5:5 ratio. A range of cytokine mix doses was tested (ESM table 1). For the 6-day experiments, the culture medium with or without cytokines was renewed every 2-3 days.

### Glucose-dependent secretion assays and hormone quantification

Islet MTs were incubated sequentially in 3D InSight Krebs Ringer HEPES Buffer (KRHB; InSphero AG, Switzerland) with 0.5% wt/vol bovine serum albumin (Sigma) with low (2.8 mmol/l) and high (16.7 mmol/l) glucose (Sigma, USA) for 2 hours each. Before initiating a secretion assay, islet MTs were washed twice, equilibrated for 1 hour, and washed again in 2.8 mmol/l glucose KRHB. Supernatants were collected and immediately frozen until hormone quantification. Glucagon was quantified using the V-PLEX Metabolic Panel 1 Human Kit (with only glucagon capture antibody, Meso Scale Diagnostics LLC, USA) on the MESO Quickplex SQ 120 instrument (Meso Scale Diagnostics, USA) and insulin was quantified using the Stellux Chemi Human Insulin ELISA (Alpco, USA) with the Spark 10M microplate reader (Tecan, Switzerland) according to the respective manufacturer’s instructions. Somatostatin was measured by RIA as previously described in [37] using antibody 1758. Technical replicates were pooled to achieve sufficient sample volume required for accurate measurement of somatostatin.

### Accumulated hormone or chemokine secretion measurements

Culture medium samples collected just prior to glucose-dependent secretion assays were used to determine ‘accumulated’ glucagon and chemokine secretion over 24 hours. Briefly, samples were immediately frozen upon collection and later quantified using the V-PLEX Metabolic Panel 1 Human Kit (with only glucagon capture antibody) and V-PLEX Chemokine Panel 1 Human Kit, respectively, on the MESO Quickplex SQ 120 instrument (all Meso Scale Diagnostics LLC, USA). The chemokine panel included Chemokine (C-X-C motif) ligand (CXCL) -8, -10, and C-C Motif Chemokine Ligand (CCL) -2, -3, -4, -11, -13, -17, -22, -26. Sample values below the limit of detection (LOD) were imputed with the LOD/√2 method [38].

### ATP and apoptosis measurements

In a subset of experiments, MTs were lysed after KRHB sample collection to determine ATP content using the CellTiter-Glo Luminescent Cell Viability Assay (with Protease Inhibitor Cocktail, both from Promega, USA). Luminescence was recorded using the Spark 10M microplate reader (Tecan, Switzerland). Apoptosis was determined as Caspase-3/7 activity immediately after 6-day culture using Caspase-Glo 3/7 Assay (Promega, USA). Luminescence was recorded with the Infinite M200 PRO microplate reader (Tecan, Switzerland).

### Immunofluorescence staining

Islet MTs were immunofluorescently stained for the beta and alpha-cell transcription factors (NKX6.1 and ARX, respectively) and for the three major islet hormones (insulin, glucagon, and somatostatin). Briefly, islet MTs were fixed for 15 minutes using 4% paraformaldehyde in PBS and permeabilised with 0.5% Triton X-100 in PBS. Blocking was performed for 1 hour using 5% donkey serum (DKS; Jackson ImmunoResearch, UK) for the ARX/NKX6.1 staining or 10% fetal calf serum (FCS; Bio&Sell, Germany) for the islet hormone staining. Following blocking, islet MTs were incubated overnight with primary antibodies in antibody incubation buffer (0.2% Triton X-100 in PBS with 5% DKS or 10% FCS), followed by a 4-hour secondary antibody incubation in antibody incubation buffer with DAPI nuclear stain. After both antibody incubations, islet MTs were washed with 0.2% Triton X-100 in PBS to eliminate nonspecific binding. Antibody sources, characteristics, and dilutions are listed in ESM Table 2.

### Confocal imaging and quantitative analysis

Post-staining, islet MTs were cleared using ScaleS4 (40% w/v D-(-) sorbitol, 24% w/v urea, 8% v/v glycerol, 0.2% v/v Triton X-100, and 15% v/v DMSO in MilliQ water [39]). Cleared islet MTs were imaged in Akura 384 ImagePro plates (InSphero AG, Switzerland) using the CQ1 benchtop high content analysis system (Yokogawa, Japan). Imaging was done using a 40x dry objective (NA 0.95). Images were acquired at a z-step size of 3 µm. Images were analysed with the CQ1 analysis software CellPathFinder (Yokogawa, Japan), using a customised pipeline to quantify DAPI-positive, ARX and NKX6.1 staining intensity or to quantify DAPI-positive, glucagon, insulin and somatostatin intensity.

### Arginine stimulation, incretin receptor agonist and/or inhibitor treatment

For the arginine stimulation test, 10 mmol/l L-arginine (Sigma-Aldrich, USA) was added acutely in the glucose-dependent secretion assay. Effects of the incretins [D-Ala^2^]-GIP, a stable GIP analogue, and liraglutide, a long-acting GLP-1RA (both from Tocris Bioscience, UK), alone and in combination, were investigated by adding them alongside the 6-day cytokine exposure, each at a concentration of 1 μmol/l. In a subset of these experiments, incretin treatment was extended to include a 24-hour preincubation period and also added during the glucose-dependent secretion assay. The effects of the incretins were furthermore investigated when added acutely during glucose-dependent secretion assays post 6-day cytokine exposure. Moreover, the effects of [D-Ala^2^]-GIP+liraglutide or tirzepatide (Selleck Chemicals LLC, USA) were investigated with or without selected inhibitors of intracellular signalling components; KT5720 (Protein kinase A (PKA) inhibitor, Tocris Bioscience, UK), HJC0350 (Exchange protein directly activated by cAMP 2 (EPAC2) inhibitor, Tocris Biosceince, UK), NKY80 (Adenylate cyclase (AC) inhibitor, Sigma-Aldrich, USA), Nimodipine (L-type calcium channel (LTCC) inhibitor, Sigma-Aldrich, USA), KN-93 (Calcium/calmodulin-dependent protein kinase II (CAMK2) inhibitor, Sigma-Aldrich, USA) or vehicle control (equivalent dilution of 0.1% DMSO). Inhibitors were added during the equilibration step and during the glucose-dependent secretion assay. Inhibitor doses were based on literature and indicated in figure legends.

### Statistical analyses

Statistical analyses of data were performed using GraphPad Prism (v10, USA). Data are presented as mean ± standard deviation (SD) for technical replicates (one donor per graph, table 1) unless otherwise stated. Outliers were identified and omitted using the robust regression and outlier removal test (ROUT), with the Q-cutoff set to 5%. Comparisons between two sample groups were analysed with unpaired Student’s *t*-tests, whereas comparisons between more than two sample groups were analysed by unpaired one-way or two-way ANOVA with Dunnett’s, Tukey’s, or Šídák’s multiple comparison tests as indicated in figure legends. Statistical significance was defined as a *p* value < 0.05, (*^/†^*p* value < 0.05, **^/††^*p* value < 0.01, ***^/†††^*p* value < 0.001), and non-significant unless stated.

## Results

### Short-term cytokine exposure increases glucagon secretion

Untreated control islet MTs exhibited a similar drop in glucagon secretion between low and high glucose glucagon secretion (∼79%) when compared to that seen in native isolated islets [40]. Using a combination of canonical type 1 diabetes cytokines, IL-1β, IFN-γ, and TNF-α, we first subjected human islet MTs from two individual donors to a short-term (1 day) cytokine exposure followed by sequential glucose-dependent secretion assays (Fig. 1a). Cytokine exposure caused a dose-dependent increase in glucagon secretion compared to untreated control MTs at both low (2.8 mmol/l) and high (16.7 mmol/l) glucose in two human islet MT donors (Fig. 1b-c). In contrast, insulin secretion was mainly affected at low glucose upon cytokine exposure (ESM Fig. 1). These results show that short-term cytokine exposure leads to aberrant glucagon and insulin secretion, but not in a pattern that resembles type 1 diabetes, i.e., reduced secretion at low and high glucose, respectively.

**Fig. 1.**
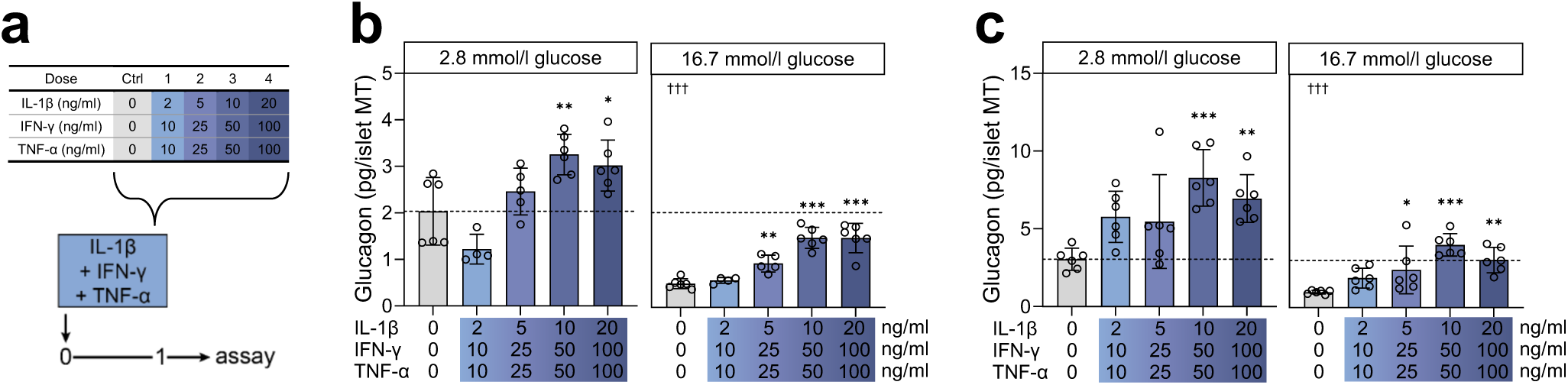
Short-term cytokine exposure increases glucagon secretion. (a) Experimental schematic of the short-term (1 day) setup. (b-c) Glucagon secretion at 2.8 and 16.7 mmol/l glucose in control (grey bars) and with increasing load of short-term cytokine exposure (blue bars) in (b) donor 1 and (c) donor 2. The dashed line denotes the baseline physiological response to low glucose of untreated control islet MTs. Data presented as mean ± SD of a single donor in 6 technical replicates. Asterisks (*) indicate statistical significance using one-way ANOVA with Dunnett’s multiple comparisons test compared to untreated control, *p <0.05, **p <0.01, ***p <0.001. Daggers (†) indicate statistical significance using Student’s t-test comparing the two untreated controls at 2.8 vs. 16.7 mmol/l glucose, †††p <0.001.

### Long-term cytokine exposure impairs glucose-dependent glucagon secretion

We next extended cytokine exposure to 6 days (long-term) before assessing the secretory function (Fig. 2a). Using islet MTs from three individual donors, we found that long-term cytokine exposure significantly diminished glucagon secretion in response to low glucose for all tested cytokine doses (Fig. 2b, d, f). Furthermore, accumulated glucagon secretion during the last 24 hours of cytokine exposure was also drastically decreased (Fig. 2c, 2e). Baseline glucagon secretion and cytokine sensitivity were highly donor-dependent. The cytokine-induced impairment of low glucose-induced glucagon secretion ranged from 15.5-43.0% (fig. 2b), 71.9-89.2% (fig. 2d), and 86.1-88.0% (fig. 2f). In islet MTs from one donor, the 6-day cytokine exposure exacerbated glucagon secretion at high glucose (92.0-146.4%, Fig. 2b), whereas it was unaffected or reduced in two donors (Fig. 2d and f). The detrimental effects of long-term cytokine exposure on high-glucose-induced insulin secretion were more consistent among the three donors, in which reductions ranged from 28.8-89.3%, 20-63.7%, and 24.4%-92.1%, respectively (ESM Fig. 2a-c). Somatostatin secretion was generally decreased upon cytokine exposure (ESM Fig. 2d-g), making paracrine inhibition of glucagon and insulin secretion by somatostatin an unlikely explanation. Collectively, these data demonstrate that long-term cytokine exposure impairs alpha-cell function during low glucose, similar to that observed in type 1 diabetes [3].

**Fig. 2.**
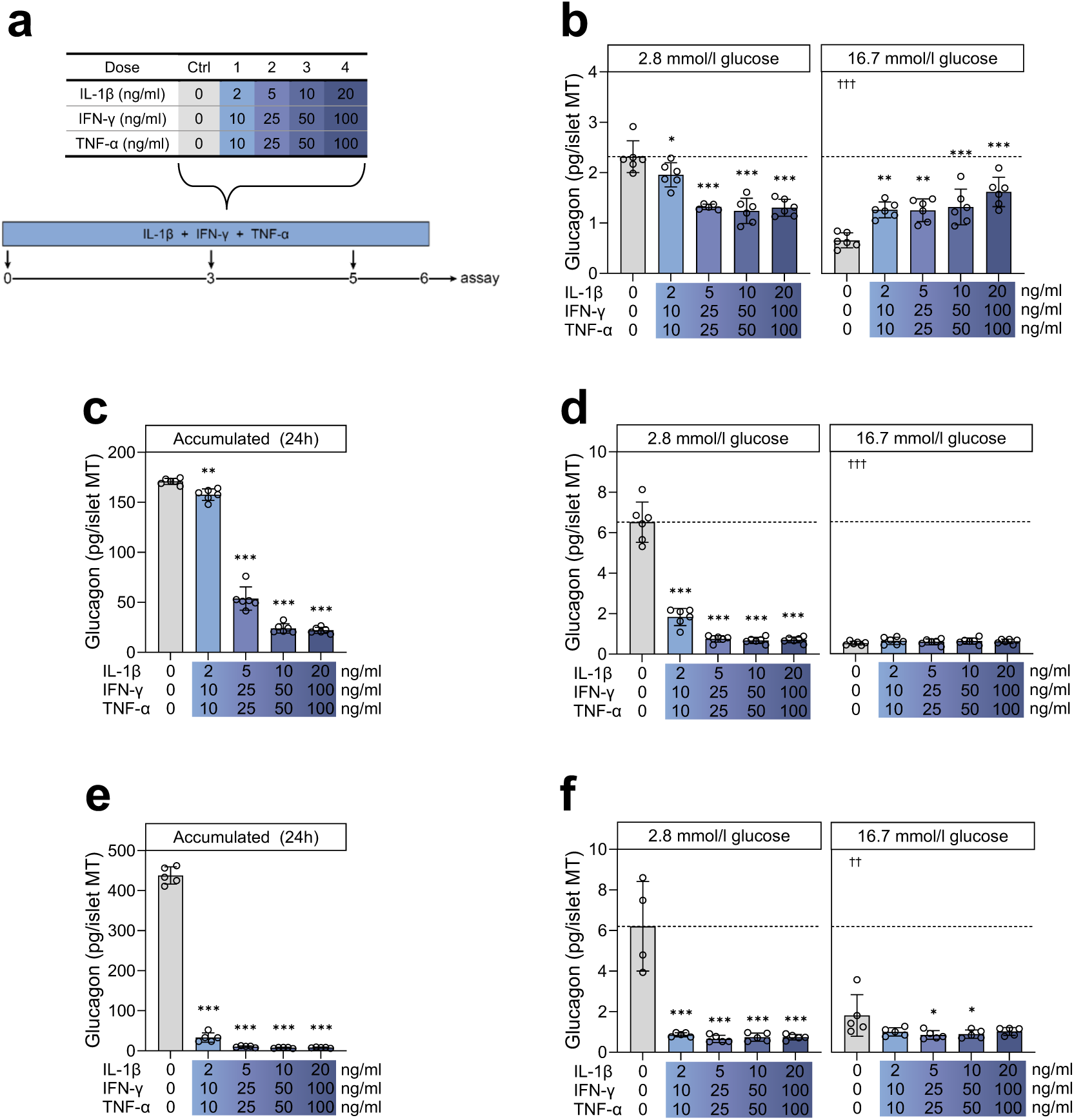
Long-term cytokine exposure impairs glucose-dependent glucagon secretion. (a) Experimental schematic of the long-term (6-day) setup. (b, d, f) Glucagon secretion at 2.8 and 16.7 mmol/l glucose in control (grey bars) and with increasing load of long-term cytokine exposure (blue bars) in (b) donor 3, (d) donor 4, and (f) donor 5. The dashed line denotes the baseline physiological response to low glucose of untreated control islet MTs. (c, e) Accumulated glucagon secretion over the last 24 hours of long-term cytokine exposure for (c) donor 4 and (e) donor 5. The dashed line denotes the untreated control baseline. Data presented as mean ± SD of a single donor in 6 technical replicates (5 technical replicates for donor 5). Asterisks (*) indicate statistical significance using one-way ANOVA with Dunnett’s multiple comparisons test compared to untreated control, *p <0.05, **p <0.01, ***p <0.001. Daggers (†) indicate statistical significance using Student’s t-test comparing the two untreated controls at 2.8 vs. 16.7 mmol/l glucose, ††p <0.01, †††p <0.001.

### Long-term cytokine exposure induces cell death, secretion of chemokines, and reduces ARX and NKX6.1, and islet hormone expression

To further support long-term cytokine exposure as a valid model of alpha-cell impairment in type 1 diabetes, we assessed islet viability using three complementary approaches: ATP content (Fig. 3a-c), caspase 3/7-activity (Fig. 3d), and confocal imaging (DAPI count, Fig. 3g, -j). In islet MTs from three donors, ATP content dropped dose-dependently in response to cytokine exposure (Fig. 3a-c). Long-term cytokine exposure also induced secretion of chemokines, i.e., immunomodulatory signalling molecules (ESM Fig. 3), such as CXCL8 and CXCL10 (ESM Fig. 3a, b). Apoptosis was increased, as reflected by an almost 10-fold increase in caspase activity after cytokine exposure of islet MTs from one donor (Fig. 3d). In line with this, cytokine-treated islet MTs from two donors showed dose-dependent reductions in DAPI-positive cell count (Fig. 3g, -j, respectively). The alpha cell identity marker ARX and beta cell identity marker NKX6.1 were dose-dependently reduced in response to cytokines (Fig. 3h-i) in one donor. Likewise, all three islet hormones (Fig. 3k-m) were dose-dependently reduced in response to cytokines in another donor.

**Fig. 3.**
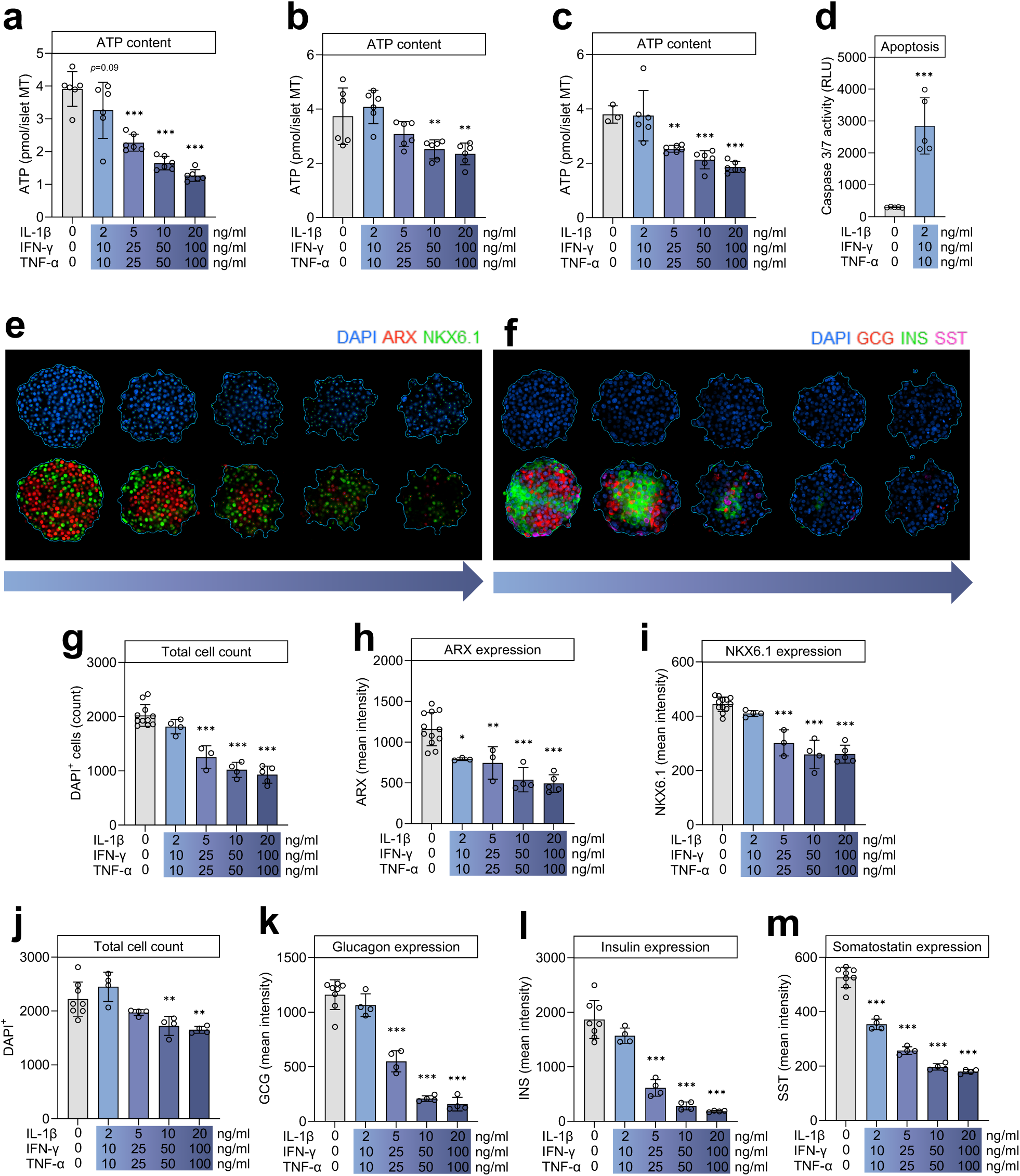
Long-term cytokine exposure induces cell death and reduces ARX, NKX6.1, and islet hormone expression. (a-c) ATP content in control (grey bars) and with increasing load of long-term cytokine exposure (blue bars) in (a) donor 3, (b) donor 4, and (c) donor 5. (d) Caspase 3/7 activity after long-term cytokine exposure (dose-¼, donor 6). (e-m) transcription factor and islet hormone stainings. (e) Representative images for transcription factor staining in islet MTs from donor 9 with increasing load of long-term cytokine exposure; DAPI (blue), ARX (red), and NKX6.1 (green) expression. (f) Representative images for islet hormone staining in islet MTs from donor X with increasing load of long-term cytokine exposure with DAPI (blue), glucagon (red), insulin (green), and somatostatin (magenta) expression. (g-i) transcription factor quantification and (j-m) islet hormone quantification. (g) DAPI-positive cell count, mean (h) ARX intensity, and (i) NKX6.1 intensity in donor 9. (j) DAPI-positive cell count, mean (k) glucagon intensity, (l) insulin intensity, and (m) somatostatin intensity in donor 10. In (a-c), data are presented as mean ± SD of a single donor in 6 technical replicates. Asterisks (*) indicate statistical significance using one-way ANOVA with Dunnett’s multiple comparisons test compared to untreated control, **p <0.01, ***p <0.001. In (d). data are presented as mean ± SD of a single donor in 5 technical replicates. Asterisks (*) indicate statistical significance using Student’s t-test comparing the untreated control vs. cytokine exposed. In (g-m), data are presented as mean ± SD of a single donor in 3-6 technical replicates (8-12 for control). Asterisks (*) indicate statistical significance using one-way ANOVA with Dunnett’s multiple comparisons test compared to untreated control, *p <0.05, **p <0.01, ***p <0.001. RLU; relative light units.

### Alpha cells partially retain secretory capacity after long-term cytokine exposure

We next examined if the alpha cells in islet MTs retain capacity to secrete glucagon in response to L-arginine following long-term cytokine exposure, as seen in type 1 diabetes [3]. For these experiments, the cytokine range was adjusted to include lower doses to optimise the window of therapeutic intervention (ESM Fig. 4 and ESM Table 1). In untreated control islet MTs, L-arginine induced a 1355% increase in glucagon secretion at low glucose (Fig. 4, *p*<0.001). For all cytokine-exposed islet MTs except those exposed to cytokine dose-2, glucagon secretion in response to L-arginine surpassed the physiological response to low glucose alone (dotted line, Fig. 4). And for the two lowest cytokine doses (dose-¼ and dose-½), L-arginine significantly induced glucagon secretion compared to cytokine-exposed islet MT without L-arginine. L-arginine also significantly increased glucagon secretion at high glucose, but to a much lesser degree than observed at low glucose. These results demonstrate that alpha cells partially retain the ability to secrete glucagon in response to L-arginine after long-term cytokine exposure.

**Fig. 4.**
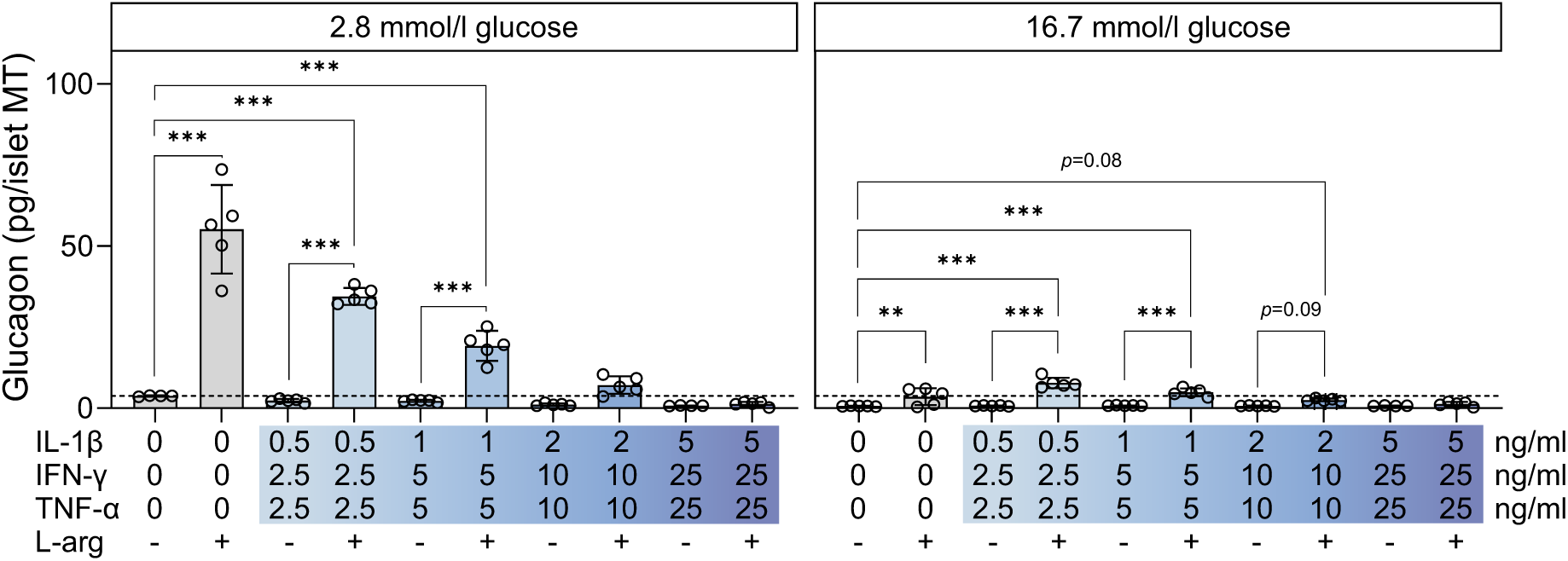
Alpha cells partially retain secretory capacity after long-term cytokine exposure. (a) Glucagon secretion at 2.8 and 16.7 mmol/l glucose in control (grey bars) and with increasing load of long-term cytokine exposure (blue bars) in donor 8 with and without (+/-) 10 mmol/l L-arginine. The dashed line denotes the baseline physiological response to low glucose of untreated control islet MTs. Data presented as mean ± SD of 5 technical replicates. Asterisks (*) indicate statistical significance using two-way ANOVA with Šídák’s multiple comparisons test comparing L-arginine stimulated (+) vs their respective L-arginine unstimulated (-) control and comparing L-arginine stimulated (+) vs the untreated control/baseline response, **p <0.01, ***p <0.001.

### Treatment with GIP and/or GLP-1 receptor agonists alongside cytokine exposure does not prevent cytokine-induced detrimental effects

Incretin receptor agonists exert anti-inflammatory and cytoprotective effects on human islets [41, 42], and liraglutide improves beta-cell function of islet MTs under cytokine stress conditions [15]. We therefore investigated if treatment with GIP and/or GLP-1 receptor agonists during long-term low-dose cytokine exposure would prevent the observed detrimental effects on glucose-dependent alpha-cell function (Fig. 5a). Long-term treatment with the stable GIP analogue [D-Ala^2^]-GIP, the long-acting GLP-1 receptor agonist liraglutide, or the combination of the two did not affect low-glucose-induced glucagon secretion either in control (Fig. 5b) or cytokine-exposed islet MTs (Fig. 5c). The inclusion of a 24-hour [D-Ala^2^]-GIP±liraglutide pre-treatment step before cytokine exposure as well as the inclusion of these during the glucose-dependent secretion assay (Fig. 5d) still failed to alter glucagon secretion (Fig. 5e-f). Long-term treatment with incretin receptor agonists alongside cytokine exposure somewhat surprisingly also failed to prevent cytokine-induced caspase 3/7 activity (Fig. 5h). As cytokines are known to stimulate islets to secrete chemokines [43], we examined the effect of [D-Ala^2^]-GIP±liraglutide on chemokine secretion during the last 24 hours of long-term exposure. While the secretion of chemokines was induced by cytokine exposure (compared to baseline levels, ESM Fig. 5), only two cytokine-induced chemokines were modulated by [D-Ala^2^]-GIP±liraglutide treatment (Fig. 6). Specifically, [D-Ala^2^]-GIP±liraglutide significantly increased CCL11 secretion (Fig. 6f; 219.8%, *p*=0.02 and 192.3%, *p*=0.04, respectively) while decreasing CCL17 secretion (Fig. 6h; 35.1%, *p*=0.05 and 38.4% *p*=0.03, respectively). The obtained data indicate that at the tested doses, treatment with incretin receptor agonists alongside cytokine exposure fails to prevent the detrimental effects of cytokines on glucagon secretion, cell death, and secretion of chemokines.

**Fig. 5.**
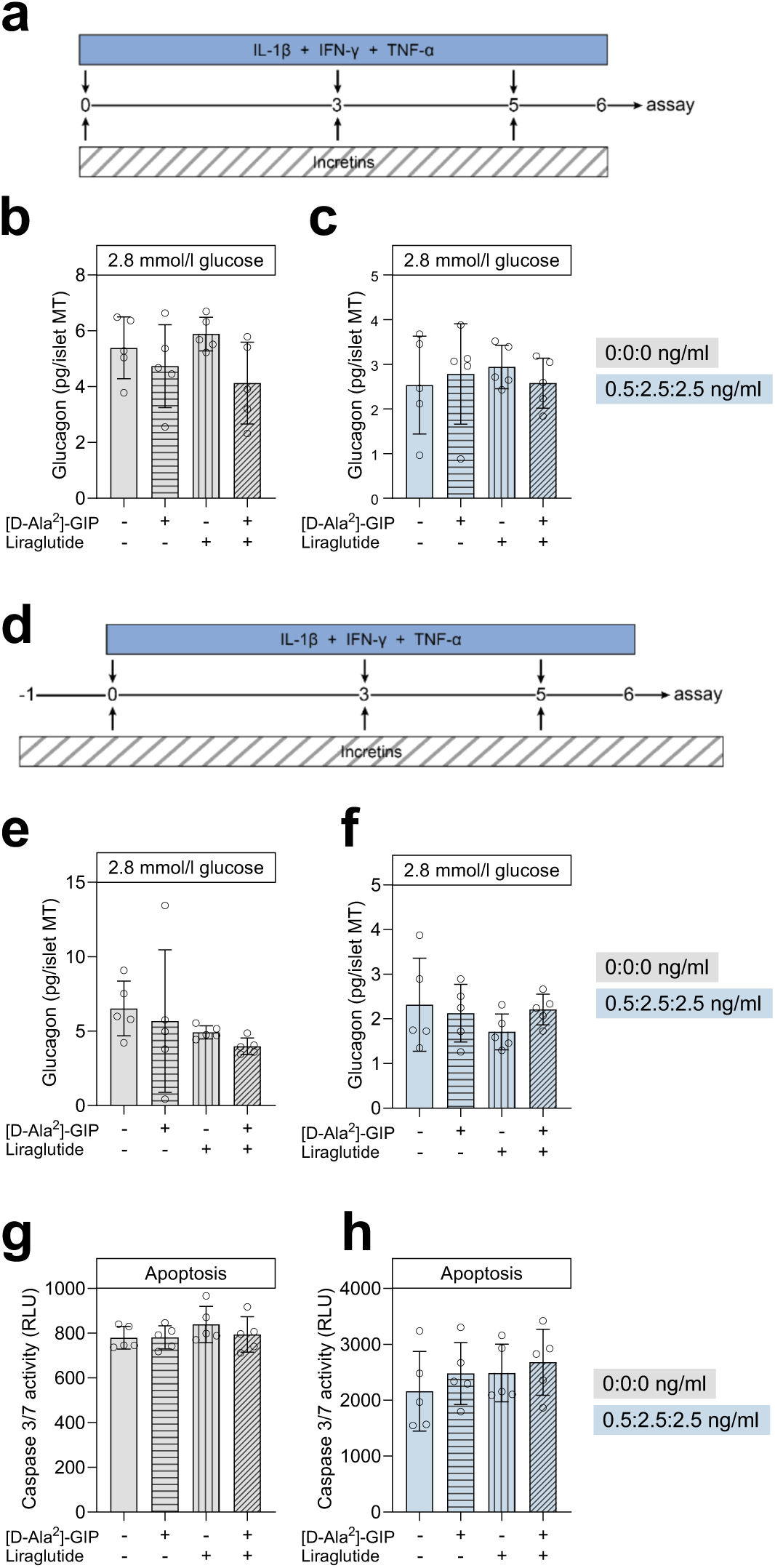
Treatment with [D-Ala^2^]-GIP ± liraglutide alongside cytokine exposure does not prevent alpha-cell impairment or cell death. (a) Experimental schematic for donor 8 (b-c). (b) Glucagon secretion at 2.8 mmol/l glucose in control (grey bars), and (c) long-term cytokine exposed (dose-¼, light blue bars) without (-/-, no bar pattern) or with 1 µmol/l [D-Ala^2^]-GIP (+/-, horizontal stripes), liraglutide (-/+, vertical stripes), or both (+/+, angled stripes). (d) Experimental schematic for donor 11 denoting addition of 24-hour pre-treatment with [D-Ala^2^]-GIP and/or liraglutide before cytokine exposure (e-h) and the addition during glucose-dependent hormone secretion assay (e-f). (e-f) same as in (b-c) but according to the schematic in (d). (g) Caspase 3/7 activity in control (grey bars), and (h) after long-term cytokine exposure (dose-¼, light blue bars) without (-/-, no bar pattern) or with 1 µmol/l [D-Ala^2^]-GIP (+/-, horizontal stripes), liraglutide (-/+, vertical stripes), or both (+/+, angled stripes). Data presented as mean ± SD of a single donor in 5 technical replicates. Results are ns using one-way ANOVA with Tukey’s multiple comparisons test. RLU, relative light units.

**Fig. 6.**
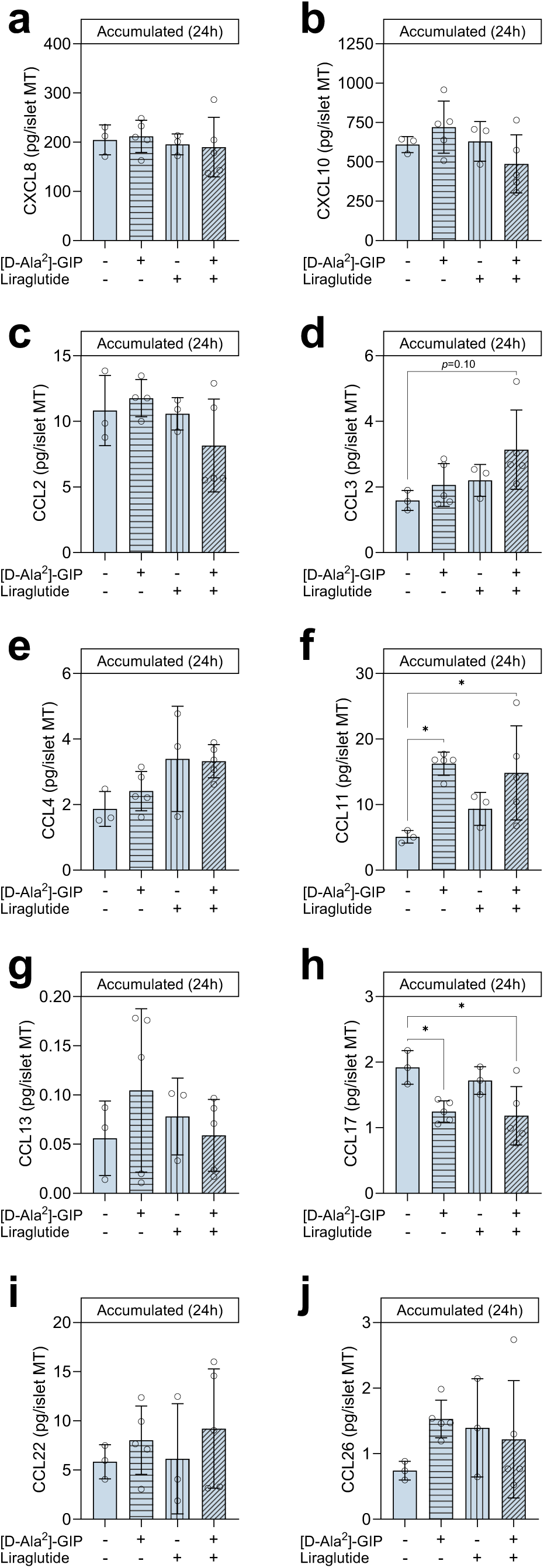
Treatment with [D-Ala^2^]-GIP ± liraglutide alters cytokine-induced secreion of CCL11 and CCL17. (a) Accumulated CXCL8, (b) CXCL10, (c) CCL2, (d) CCL3, (e) CCL4, (f) CCL11, (g) CCL13, (h) CCL17, (i) CCL22, and (j) CCL26 secretion during the last 24 hours of the long-term cytokine exposure (dose-¼) without (-/-, no bar pattern) or with 1 µmol/l [D-Ala^2^]-GIP (+/-, horizontal stripes), liraglutide (-/+, vertical stripes), or both (+/+, angled stripes). Data presented as mean ± SD of donor 11 in 5 technical replicates (only 3 technical replicates for untreated control and liraglutide bars due to pipetting error). Asterisks (*) indicate statistical significance using one-way ANOVA with Tukey’s multiple comparisons test, *p <0.05.

### Acute treatment with GIP and/or GLP-1 receptor agonists after cytokine exposure partially restores glucose-dependent glucagon secretion

Next, we tested if [D-Ala^2^]-GIP±liraglutide treatment *after* long-term cytokine exposure would alleviate the cytokine-induced detrimental effects on glucagon secretion (Fig. 7a). Acute treatment with [D-Ala^2^]-GIP+liraglutide following cytokine exposure yielded a 118.8% increase in baseline low-glucose-induced glucagon secretion (Fig. 7b, *p*<0.001). A comparable increase was observed for islet MTs exposed to cytokine dose ¼ (139.5%, *p*<0.001), in which alleviation even surpassed the baseline glucagon secretion response to low glucose (dashed line, Fig. 7b). At higher cytokine doses, [D-Ala^2^]-GIP+liraglutide could no longer significantly boost glucagon secretion although the glucagon levels were higher for all incretin receptor agonist-treated islet MTs (by 34.5%, 90.1%, and 75.2%).

**Fig. 7.**
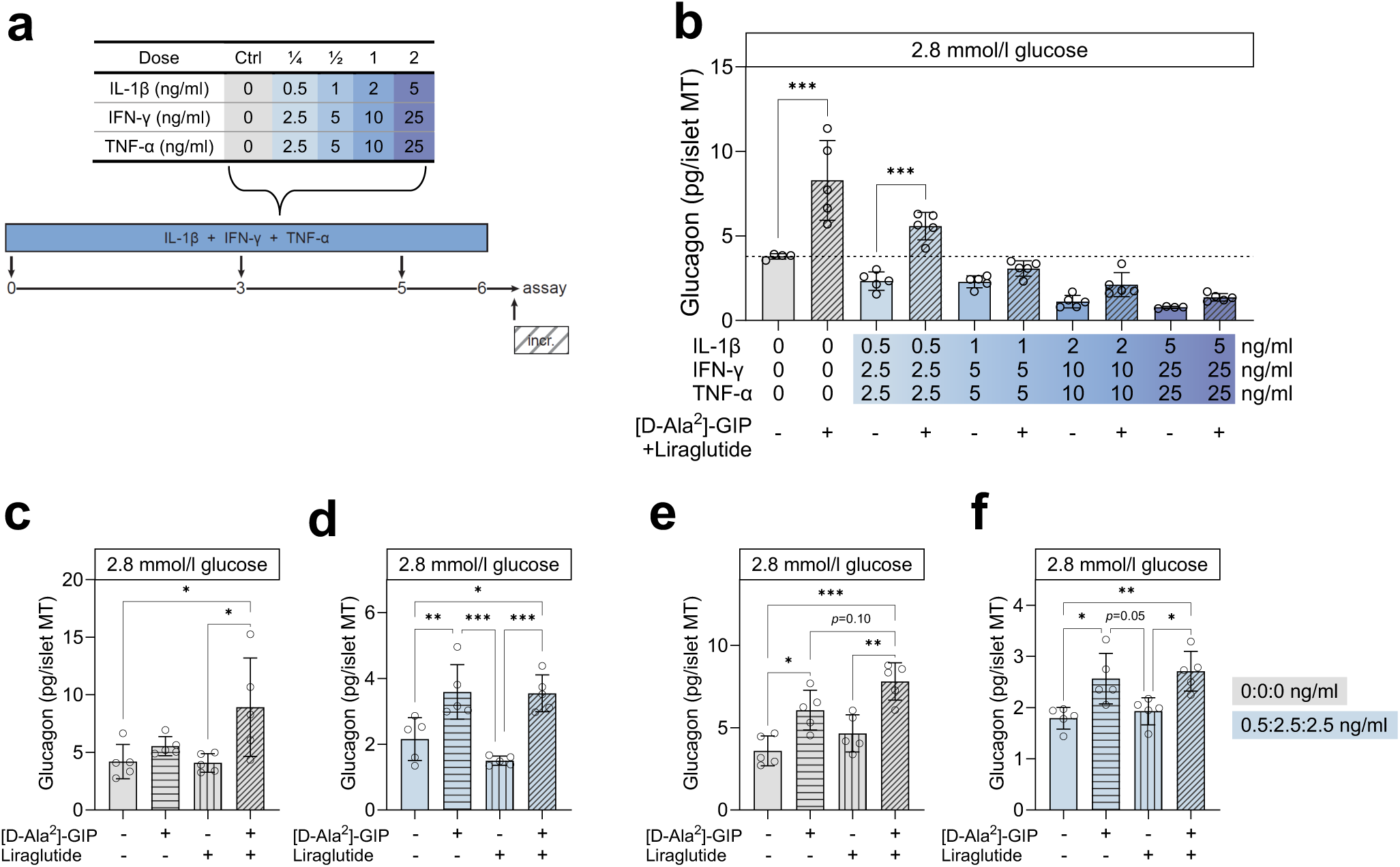
Acute treatment with [D-Ala^2^]-GIP ± liraglutide following long-term cytokine exposure boosts glucose-dependent glucagon secretion. (a) Experimental schematic denoting acute [D-Ala^2^]-GIP and/or liraglutide treatment during hormone secretion assay. (b) Glucagon secretion at 2.8 mmol/l glucose in control (grey bars) or with increasing load of long-term cytokine exposure (blue bars) in donor 8. Bar pattern (angled stripes) and (-/+) denote acute treatment with [D-Ala^2^]-GIP+liraglutide (1 µmol/l each) during the hormone secretion assay. (c) Glucagon secretion at 2.8 mmol/l glucose for donor 11 in control (grey bars), and (d) cytokine-exposed islet MTs (dose-¼, light blue bars) without (-/-, no bar pattern) or followed by acute treatment with 1 µmol/l [D-Ala^2^]-GIP (+/-, horizontal stripes), liraglutide (-/+, vertical stripes), or both (+/+, angled stripes). (e-f) same as (c-d) but for donor 7. Data presented as mean ± SD of a single donor in 5 technical replicates. In (b), asterisks (*) indicate statistical significance using two-way ANOVA with Šídák’s multiple comparisons test comparing [D-Ala^2^]-GIP+liraglutide treated vs their respective untreated control. In (c-f), asterisks (*) indicate statistical significance using one-way ANOVA with Tukey’s multiple comparisons test, *p <0.05, **p <0.01, ***p <0.001.

We next examined the individual contribution of [D-Ala^2^]-GIP and liraglutide to the partial restoration of cytokine-induced alpha-cell impairment (Fig. 7c-f). [D-Ala^2^]-GIP alone or in combination with liraglutide yielded similar increases in glucagon secretion in two donors: 66.5% (*p*=0.008) and 64.5% (*p*=0.01) in donor 8 (Fig. 7d) and 43.1% (*p*=0.02) and 51.2% (*p*=0.004) in donor 9 (Fig. 7f). In untreated control islet MTs, liraglutide tended to potentiate [D-Ala^2^]-GIP-induced glucagon secretion; 32.0% vs. 2.1 112.7% (*p*=0.14, Fig. 7c) and 69% vs. 117.5% (*p*=0.10, Fig. 7e). Liraglutide alone did not significantly affect glucagon secretion.

### Small-molecule inhibitors of calcium and cyclic AMP signalling components do not abrogate incretin-induced glucagon secretion

To gain insight into the signalling factors involved in the incretin receptor agonist-induced alleviation of cytokine-induced alpha-cell impairment, we used small-molecule inhibitors of PKA, AC, LTCC, EPAC2, and CaMK2 to see if any of these would counteract the [D-Ala^2^]-GIP±liraglutide-induced effect. In these experiments, we included the single-molecule dual GIPR and GLP-1R agonist tirzepatide alongside [D-Ala^2^]-GIP+liraglutide (Fig. 8). Similar to what was observed for [D-Ala^2^]-GIP+liraglutide, tirzepatide treatment augmented glucagon secretion in both untreated control (Fig. 8a, -c) and cytokine-exposed islet MTs (Fig. 8b, -d). None of the tested inhibitors (KT5720, HJC0350, NKY80, Nimodipine, and KN-93) reduced baseline, GIP+liraglutide-induced or tirzepatide-induced glucagon secretion of untreated control (8a, -c) or cytokine-exposed islet MTs (fig. 8b, -d), suggesting that AC, PKA, EPAC2, LTCC, and CaMK2 are not required or are possibly redundant in mediating the dual incretin receptor agonist-induced effect on alpha-cell function in islet MTs.

**Fig. 8.**
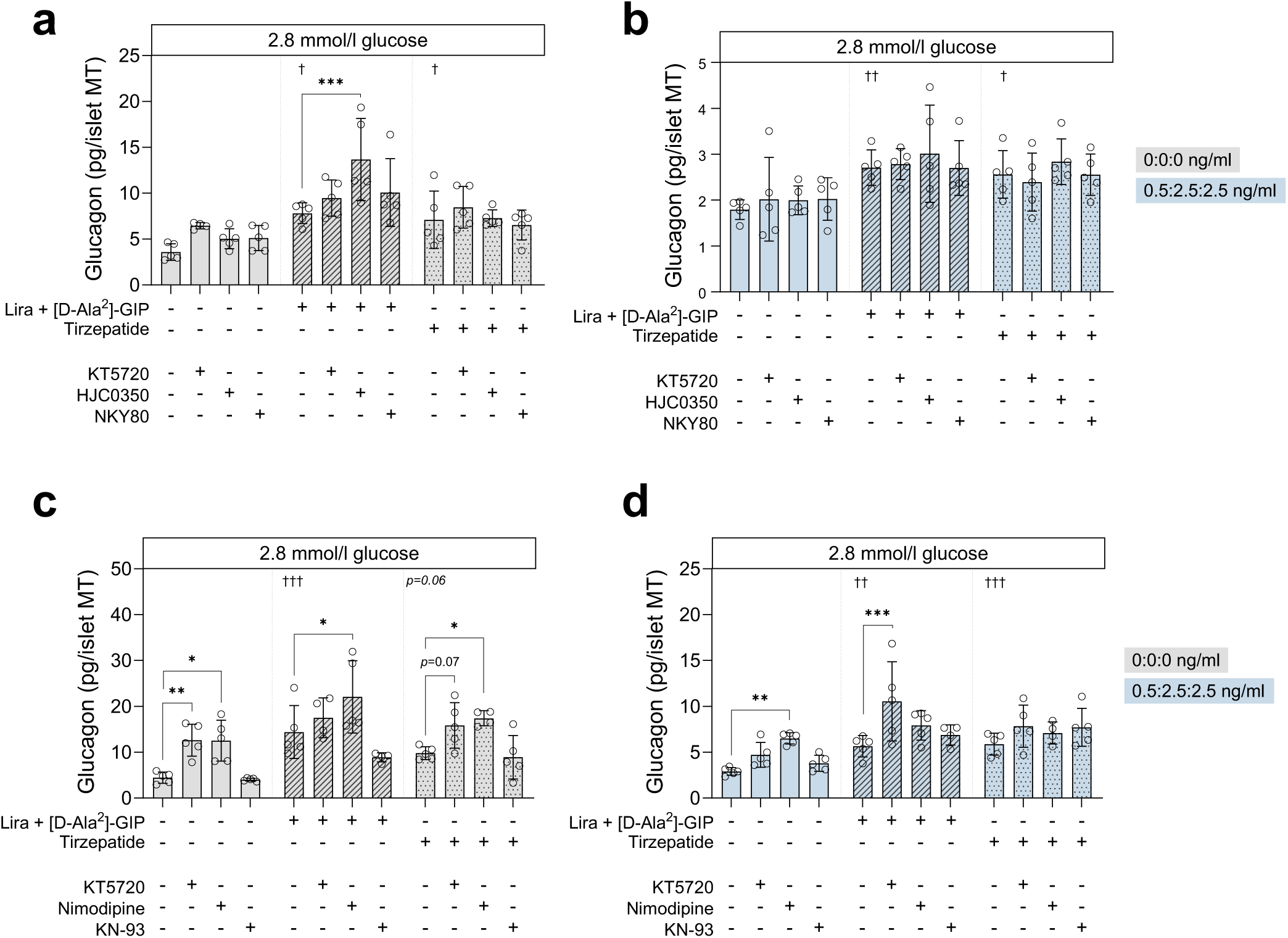
Tirzepatide- or D-[Ala^2^]-GIP+liraglutide-induced glucagon secretion is not reduced by inhibitors of cAMP or Ca^2+^ signalling components. (a) Glucagon secretion at 2.8 mmol/l glucose in control (grey bars) and (b) cytokine-exposed islet MTs (dose-¼, light blue bars) without (no bar pattern) or with acute [D-Ala^2^]-GIP+liraglutide (1 µmol/l each, angled stripes) or 1 µmol/l tirzepatide (dotted grid) treatment ± KT5720 (PKAi, 1 μmol/l), HJC0250 (EPAC2i, 5 μmol/l) or NKY80 (ACi, 20 μmol/l) inhibitors as indicated by (+) in donor 7. (c-d) same as (a-b) but for donor 12 using KT5720 (PKAi, 1 μmol/l), nimodipine (LTCCi, 10 μmol/l) and KN-93 (CaMK2i, 10 μmol/l) inhibitors. Data presented as mean ± SD of a single donor in 5 technical replicates. Asterisks (*) indicate statistical significance using two-way ANOVA with Dunnett’s multiple comparisons test, *p <0.05, **p <0.01, ***p <0.001. Daggers (†) indicate statistical significance using one-way ANOVA comparing [D-Ala^2^]-GIP+liraglutide or tirzepatide to the untreated control without inhibitors, †p <0.05, ††p <0.01, †††p <0.001.

## Discussion

In this study, we characterised alpha-cell sensitivity to diabetogenic cytokines to establish a valid in vitro model of alpha-cell impairment relevant to type 1 diabetes using islet MTs. Our data demonstrate that long-term cytokine exposure leads to alpha-cell impairment that can be partly restored by treatment with [D-Ala^2^]-GIP with or without liraglutide or by tirzepatide.

Short-term cytokine exposure increased glucagon secretion dose-dependently at both low and high glucose. Although post-prandial (i.e., high glucose) glucagon hypersecretion has been described early in type 1 diabetes disease progression [44], short-term cytokine exposure did not appear to reflect a type 1 diabetes phenotype with regards to the hypoglycaemic state (low glucose), characterised by reduced glucagon secretion [3–6]. This increase in glucagon secretion may reflect an initial response to cytokines, as increased glucagon has been shown to correlate with inflammation [45]. Alternatively, the increased glucagon secretion may reflect alpha-cell dysfunction preceding beta-cell loss, as described for autoantibody-positive donors [46]. This is further supported by glucose-stimulated insulin secretion being only modestly altered at high glucose. Indeed, while reduced glucose-stimulated insulin secretion has been observed in rodent models already in the time range of 24-48 hours, human islets appear to require longer exposure times [47, 48]. This is supported by recent studies using islet MTs, which showed that prolonged cytokine challenge reduced glucose-stimulated insulin secretion [14, 15].

Our data show that prolonged (6-day) cytokine exposure culminated not only in impaired beta-cell function but also suppressed alpha- and delta-cell function. Of note, in two of the three islet MT donors tested, maximal suppression in low-glucose-induced glucagon secretion was evident at the lowest cytokine dose (dose-1), a pattern we did not observe for high-glucose-induced insulin secretion in the same donors. These findings suggest that glucagon secretion might be more sensitive to the direct effects of cytokines than insulin secretion, at least under the experimental conditions in the current study. Consistent with this notion, the immunofluorescence stainings indicated that ARX expression was affected by cytokines already at the lowest cytokine dose compared to NKX6.1, possibly suggesting an early loss of alpha-cell identity. However, it is important to note that we only performed staining of islet MTs from a single donor, and we only evaluated based on one identity marker. Additional experiments should validate these results. Several putative mechanisms behind the impaired counterregulatory glucagon secretion in type 1 diabetes have been proposed. Most likely, impairment cannot be ascribed to a single responsible factor or mechanism but the consequence of an interplay between several factors related to paracrine signalling—for instance, loss of intra-islet insulin or increased somatostatin secretion [49, 50]. Despite the strongly impaired low-glucose-induced glucagon secretion, the ability to respond to arginine was partially maintained, especially in islet MTs exposed to the lower range of cytokine doses, suggesting the presence of alpha cells with secretory capacity exceeding basal low-glucose secretion and an assay window suitable for translational exploration of augmentation of glucagon responses.

The pathophysiology of type 1 diabetes is characterised by islet inflammation [51, 52]. Both beta [43] and alpha cells [53] play an active role during type 1 diabetes progression, e.g., via secretion of chemokines and cytokines with immune-attractant effects. Consistent with this, we found the secretion of 9 well-known chemokines was upregulated upon long-term cytokine exposure. Hence, islet MTs, similar to native human islets and beta-cell models, secrete chemokines in response to cytokines. Also consistent with type 1 diabetes pathophysiology, our model demonstrated dose-dependent reductions in ATP content and increased caspase 3/7 activity indicative of apoptosis.

We and others have described GIP as a dual-acting hormone with both insulinotropic and glucagonotropic effects in isolated human islets [40, 54], in healthy individuals [29, 55], and in individuals with type 1 diabetes [31]. GLP-1RAs have been described in pre-translational beta-cell models as anti-apoptotic and to prevent loss of glucose-stimulated insulin secretion [15, 41, 42]. We hypothesised that dual incretin receptor agonism would be superior to GIP receptor agonism alone in preventing the cytokine-induced type 1 diabetes alpha-cell phenotype indirectly via preservation of beta cells and/or beta-cell function. Contrary to this hypothesis, treatment with [D-Ala^2^]-GIP+liraglutide alongside cytokine exposure did not prevent development of a type 1 diabetes alpha-cell phenotype, and liraglutide alone did not reduce apoptosis in the tested conditions. No protective effects of liraglutide in terms of reduced secretion of chemokines were observed, as only [D-Ala^2^]-GIP alone significantly changed the secretion of two chemokines (CCL11 and CCL17), though in opposite directions. The lack of effect of liraglutide could, at least in part, be explained by liraglutide’s palmitic acid chain that binds to albumin for stability, which necessitates high concentrations for in vitro experiments to have an adequate available pool of liraglutide for receptor binding [56]. Exceeding the 1 μmol/l concentration used here may enhance liraglutide impact. In the experiments with treatment alongside cytokine exposure, [D-Ala^2^]-GIP alone did not prevent the development of the type 1 diabetes phenotype of alpha cells either. However, GIPR desensitisation due to chronic GIP has been reported in mouse adipocytes [57], and human GIPR appears to be more prone to desensitisation through β-arrestin recruitment [58]. Hence, the lack of an alpha-cell-preservative effect in our study could be due to the chronic nature of [D-Ala^2^]-GIP treatment, causing desensitisation of the GIPR. In contrast, acute [D-Ala^2^]-GIP treatment augmented glucagon secretion in untreated control islet MTs and partially restored glucagon secretion in long-term cytokine-exposed islets. Interestingly, despite reports of glucagonostatic effects of GLP-1 on glucagon secretion at low glucose [59, 60], [D-Ala^2^]-GIP+liraglutide was comparable to [D-Ala^2^]-GIP in augmenting glucagon secretion. Furthermore, liraglutide alone did not inhibit glucagon secretion. GLP-1 effects on alpha cells are complex and have mainly been considered indirect, as GLP-1R expression is thought to be limited or absent in human alpha cells, although uncertainty exists [59, 61, 62]. However, there is some evidence that GLP-1 can act directly on alpha cells, as glucagonostatic effects are independent of insulin and somatostatin, as evidenced by INSR and SSTR2 blockage [59]. A higher incidence of hypoglycaemia (but not severe hypoglycaemia) has been associated with GLP-1RA adjunctive therapy [63], this could be due to a lesser need for exogenous insulin that is not promptly adjusted in treatment regimens. In any case, the glucagonostatic effect at low glucose remains controversial, as the glucagonostatic effects of GLP-1RAs have mainly been described in the setting of fasting hyperglycaemia or post-prandial hyperglycaemia [62, 64, 65]. In addition, several studies have found that GLP-1RAs did not compromise the hypoglycaemic glucagon in healthy and type 1 diabetes individuals [66, 67]. The ambiguous findings in the literature underline the complexity of incretin-islet biology, demanding further studies, especially in human islet model systems.

Importantly, we found comparable effects of [D-Ala^2^]-GIP+liraglutide and tirzepatide with regard to low-glucose-induced glucagon secretion. This observation is supported by the apparent main contribution of [D-Ala^2^]-GIP in driving the glucagonotropic effect when in combination with liraglutide – and tirzepatide has been reported as an imbalanced GIPR/GLP-1R agonist (higher receptor engagement at the GIPR than GLP-1R and biased towards cAMP generation at the GLP-1R) [35, 68]. Additionally, while the glucagonotropic effect of tirzepatide during low glucose has been established [35], we now, importantly, extend this to post-cytokine exposure.

In our attempt to elucidate the signalling factors indispensable for partially restoring glucagon secretion in islet MTs, we failed to identify any specific underlying mechanisms. The application of small-molecule inhibitors of various signalling components related to cAMP or calcium signalling, including AC, PKA, EPAC2, LTCC, or CaMK2, did not abrogate incretin receptor agonist-induced glucagon secretion in the tested conditions. The culmination of downstream signalling events requires further exploration, including comparing major islet cell types separately.

In summary, this study presents an in vitro model of alpha-cell impairment in type 1 diabetes. The model enables functional studies and screening of compounds to resurrect glucose-dependent glucagon secretion. Our study also shows for the first time that [D-Ala^2^]-GIP can ameliorate alpha-cell impairment to low glucose after cytokine exposure, which was not opposed by liraglutide. Additionally, tirzepatide, recognised as an imbalanced incretin co-agonist, can ameliorate alpha-cell impairment to low glucose following cytokine exposure. The presented findings should be considered in light of limitations inherent in human donor studies, specifically stemming from the restricted pool of donors.

## Additional Information

## Acknowledgements

The authors gratefully acknowledge organ donors and the next of kin of organ donors, without whom this research would not be possible. The authors thank Rebekka Gerwig and Lene Brus Albæk for the technical support with the chemokine and somatostatin measurements, respectively.

## Data Availability

The datasets generated and analysed during the study are available from the corresponding author upon reasonable request.

## Authors’ relationships and activities

KH has nothing to declare. CR, ACT, SJ and BY are employees of InSphero. B.H. is a co-founder of Bainan Biotech. J.J.H. has served on scientific advisory panels for and/or has received speaker honoraria from Novo Nordisk and MSD/Merck, is co-founder and board member of Antag Therapeutics, and co-founder of Bainan Biotech. FKK has served on scientific advisory panels and/or been part of speaker’s bureaus for, served as a consultant to, and/or received research support from Amgen, AstraZeneca, Bayer, Boehringer Ingelheim, Carmot Therapeutics, Eli Lilly, Gubra, MedImmune, MSD/Merck, Mundipharma, Norgine, Novo Nordisk, Sanofi, ShouTi, Zealand Pharma and Zucara. FKK is a co-founder of and minority shareholder in Antag Therapeutics, owns stocks in Eli Lilly, Novo Nordisk and Zealand Pharma, and has been employed by Novo Nordisk since 1st December 2023. JS owns stocks in Novo Nordisk A/S.

## Contribution statement

KH, FKK, and JS conceptualised the study. KH, JS, and BY designed the experiments. CR, SJ, and ACT contributed to the data acquisition and analysis. BH and JJH were responsible for the somatostatin RIA measurements. KH wrote the initial draft. All authors provided critical scientific input to the manuscript and approved the final version.

## Abbreviations

ARX: Aristaless-related homeobox
NKX6.1: NK6 homeobox 1
MT: Microtissue
GIP: Glucose-dependent insulinotropic polypeptide
GLP-1: Glucagon-like peptide 1
KRHB: Krebs ringer HEPES buffer
RA: Receptor agonist
CCL: C-C Motif Chemokine Ligand
CXCL: Chemokine (C-X-C motif) ligand
PKA: Protein kinase A
EPAC2: Exchange protein directly activated by cAMP 2
AC: Adenylate cyclase
LTCC: L-type calcium channel
CaMK2: Calcium/calmodulin-dependent protein kinase II
NS: Non-significant
RLU: Relative light units

## ESM Tables

**ESM Table 1.**
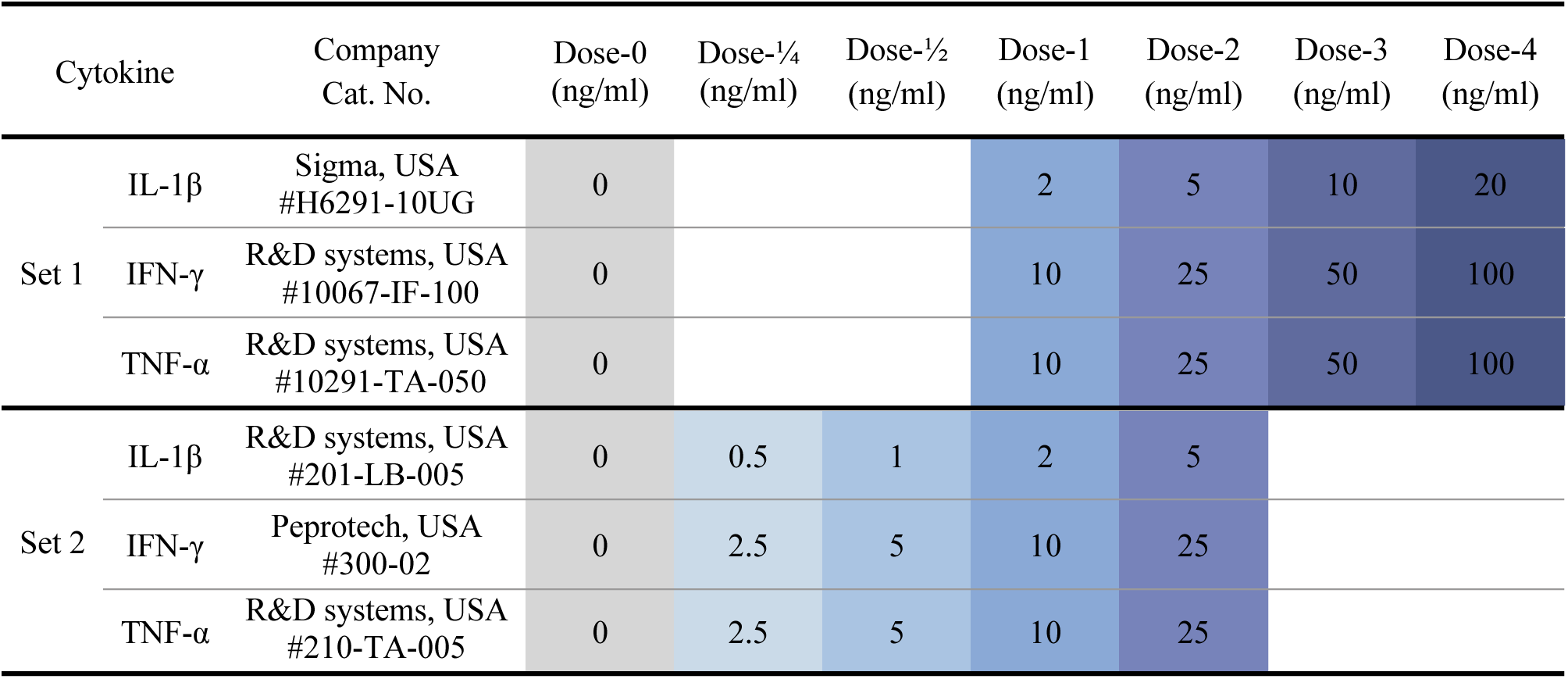
Related to Methods. Cytokines and doses.

**ESM Table 2.** Related to Methods. Antibodies and dilutions used for immunofluorescent staining and confocal imaging.

| Antibody | Supplier, #Cat. number | Dilution |
| --- | --- | --- |
| *Primary antibodies* |  |  |
| Sheep anti-ARX | R&D Systems, USA<br>(#AF7068-SP) | 1:400 |
| Rabbit anti-NKX6.1 | Abcam, UK<br>(#AB221549) | 1:200 |
| Mouse anti-Glucagon | Sigma-Aldrich, USA<br>(#G2654) | 1:300 |
| Mouse anti-Insulin (AF488) | Invitrogen, USA<br>(#53-9769-82) | 1:100 |
| Rat anti-Somatostatin | Invitrogen, USA<br>(#MA5-16987) | 1:200 |
| *Secondary antibodies* |  |  |
| Donkey anti-sheep (AF647) | Jackson ImmunoResearch, UK<br>(#713-605-147) | 1:200 |
| Donkey anti-rabbit (AF568) | Invitrogen, USA<br>(#A10042) | 1:200 |
| Alpaca anti-mouse (AF568) | ChromoTek, GER<br>(#sms1AF568-1-100) | 1:200 |
| Goat anti-rat (AF647) | Jackson ImmunoResearch, UK<br>(#112-605-167) | 1:100 |
| *Fluorescent stains* |  |  |
| DAPI | Sigma-Aldrich, USA<br>(#D9542) | 1:500 |

## ESM Figures

**ESM Fig. 1.**
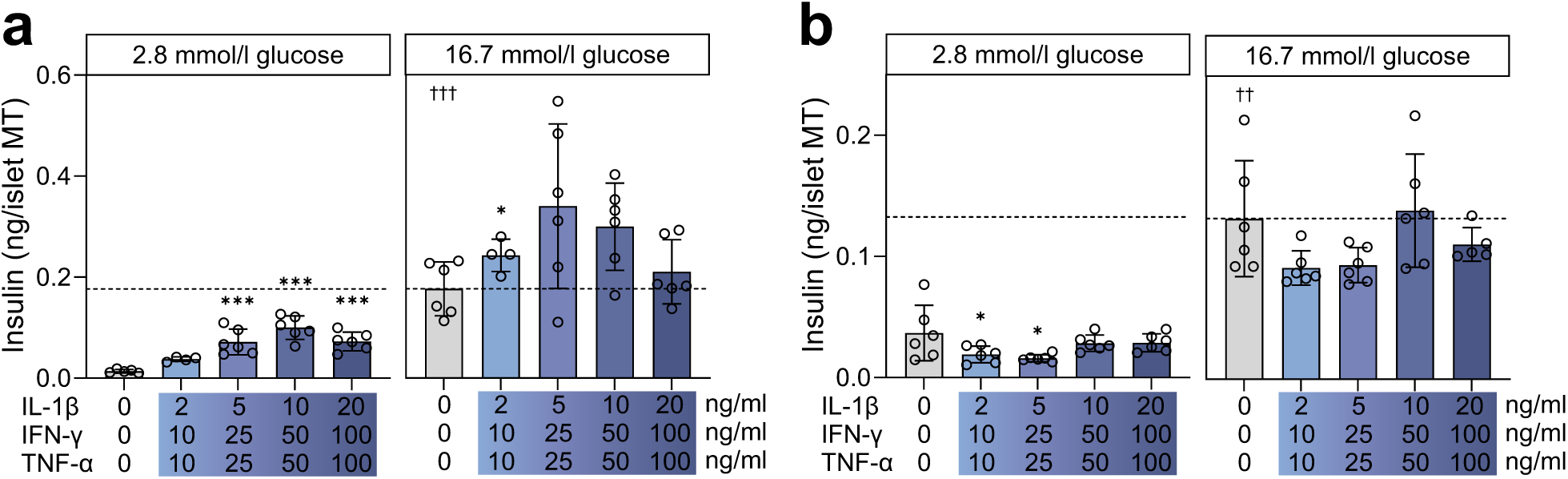
Related to Fig. 1. (a-b) Insulin secretion at 2.8 and 16.7 mmol/l glucose in control (grey bars) and with increasing load of short-term cytokine exposure (blue bars) in (a) donor 1 and (b) donor 2. The dashed line denotes the baseline physiological response to high glucose of untreated control islet MTs. Data presented as mean ± SD of a single donor in 6 technical replicates. Asterisks (*) indicate statistical significance using one-way ANOVA with Dunnett’s multiple comparisons test compared to untreated control, *p <0.05, ***p <0.001. Daggers (†) indicate statistical significance using Student’s t-test comparing the two untreated controls at 2.8 vs. 16.7 mmol/l glucose, ††p <0.01, †††p <0.001.

**ESM Fig. 2.**
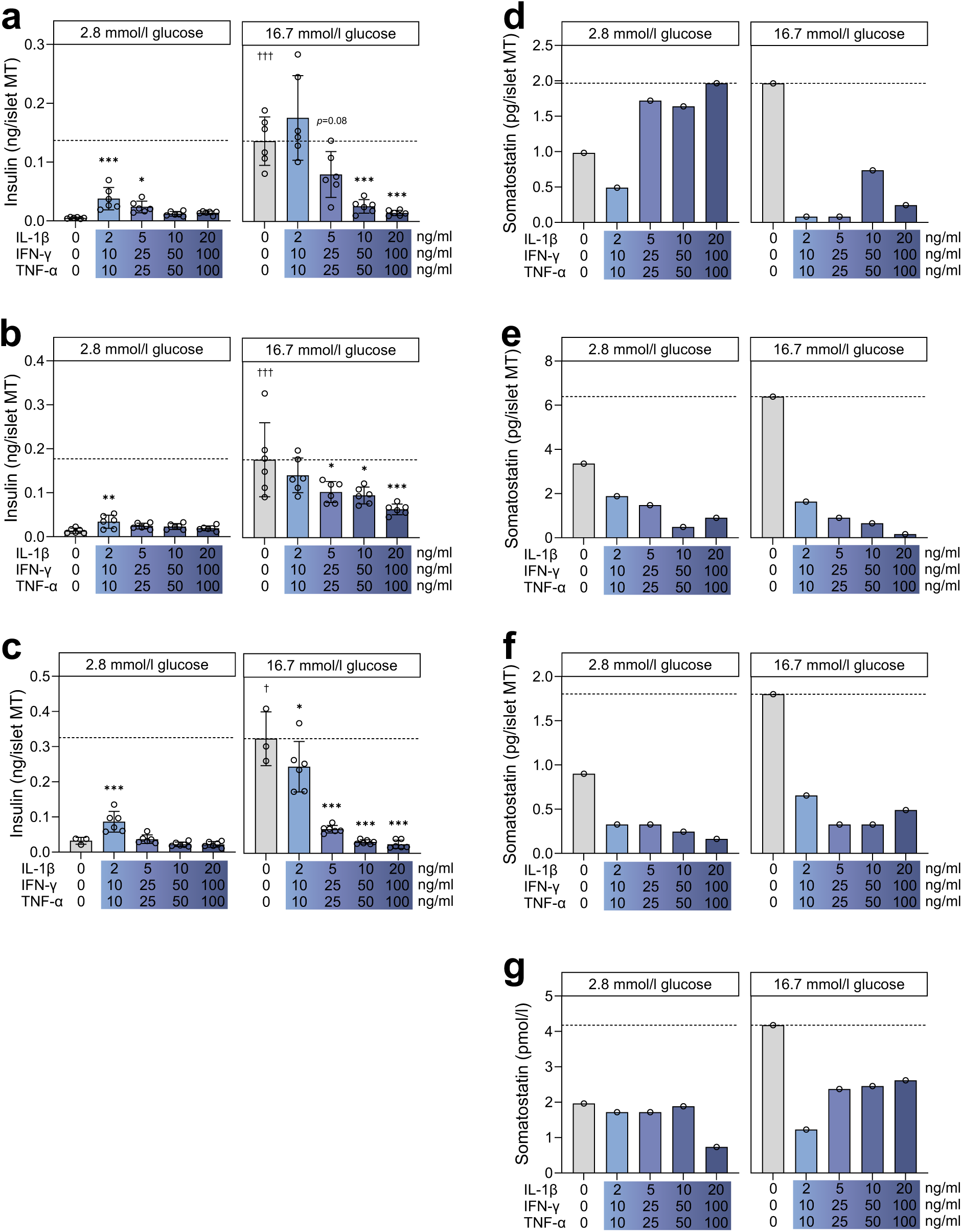
Related to Fig. 2. (a-c) Insulin secretion at 2.8 and 16.7 mmol/l glucose in control (grey bars) and with increasing load of long-term cytokine exposure (blue bars) in (a) donor 3, (b) donor 4, and (c) donor 5 islet MTs. The dashed line denotes the baseline physiological response to high glucose of untreated control. (d-g) Somatostatin secretion at 2.8 and 16.7 mmol/l glucose in control (grey bars) and with increasing load of long-term cytokine exposure (blue bars) in (d) donor 1, (e) donor 2, (f) donor 6, and (g) donor 7. In (a-c), data are presented as mean ± SD of a single donor in 6 technical replicates. Asterisks (*) indicate statistical significance using one-way ANOVA with Dunnett’s multiple comparisons test compared to untreated control, *p <0.05, **p <0.01, ***p <0.001. Daggers (†) indicate statistical significance using Student’s t-test comparing the two untreated controls at 2.8 vs. 16.7 mmol/l glucose, †p <0.05, †††p <0.001. In (d-g), data represent values measured from 6 pooled technical replicates from a single donor. No statistics were applied due to the sample size limitation.

**ESM Fig. 3.**
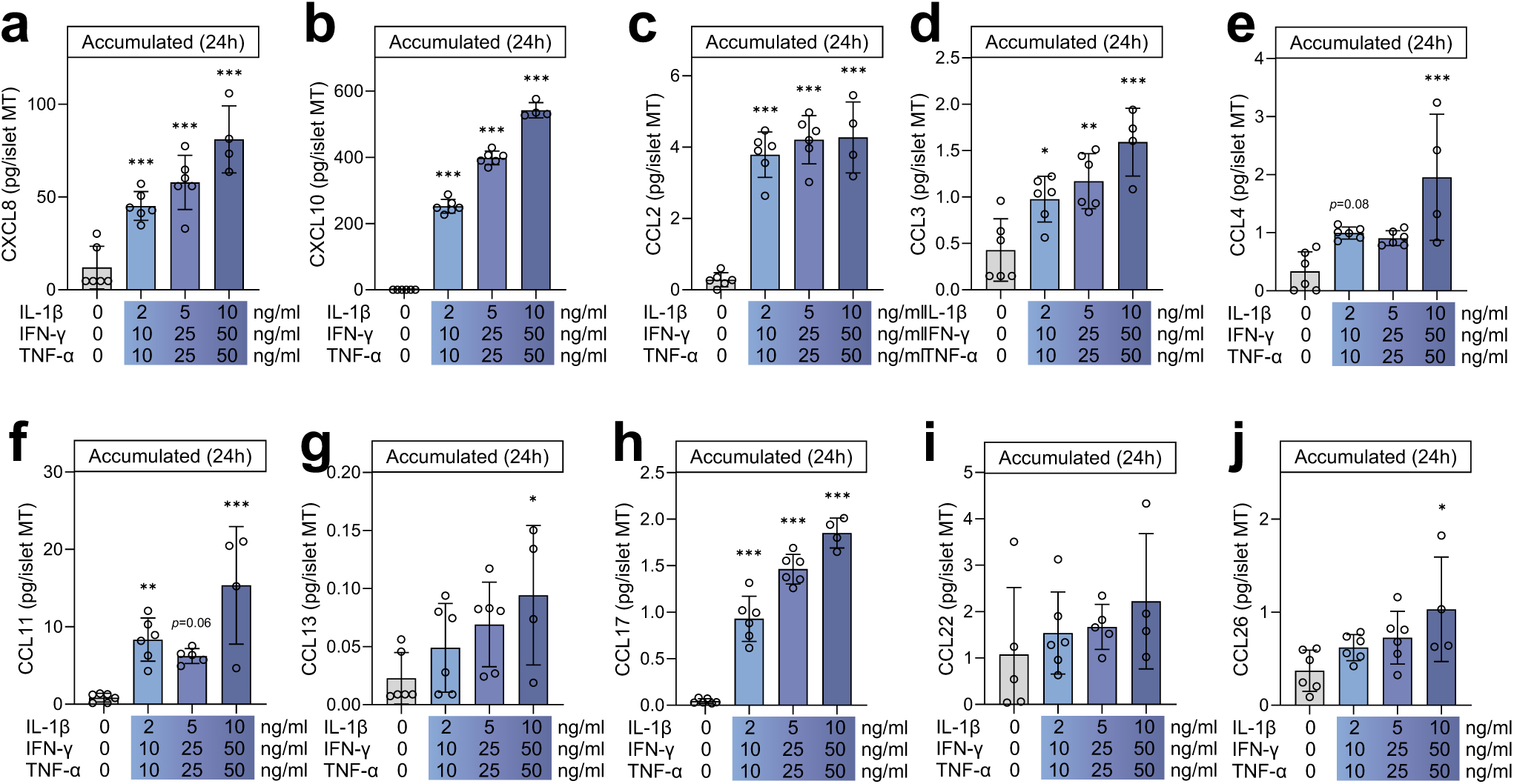
Related to Fig. 3. (a) Accumulated CXCL8, (b) CXCL10, (c) CCL2, (d) CCL3, (e) CCL4, (f) CCL11, (g) CCL13, (h) CCL17, (i) CCL22, and (j) CCL26 secretion during the last 24 hours of increasing long-term cytokine exposure (blue bars) or untreated control (grey bar) in donor 4. Data presented as mean ± SD of a single donor in 6 technical replicates (only 4 technical replicates for dose-3 due to technical error). Asterisks (*) indicate statistical significance using one-way ANOVA with Dunnett’s multiple comparisons test compared to untreated control, *p <0.05, **p <0.01, ***p <0.001.

**ESM Fig. 4.**
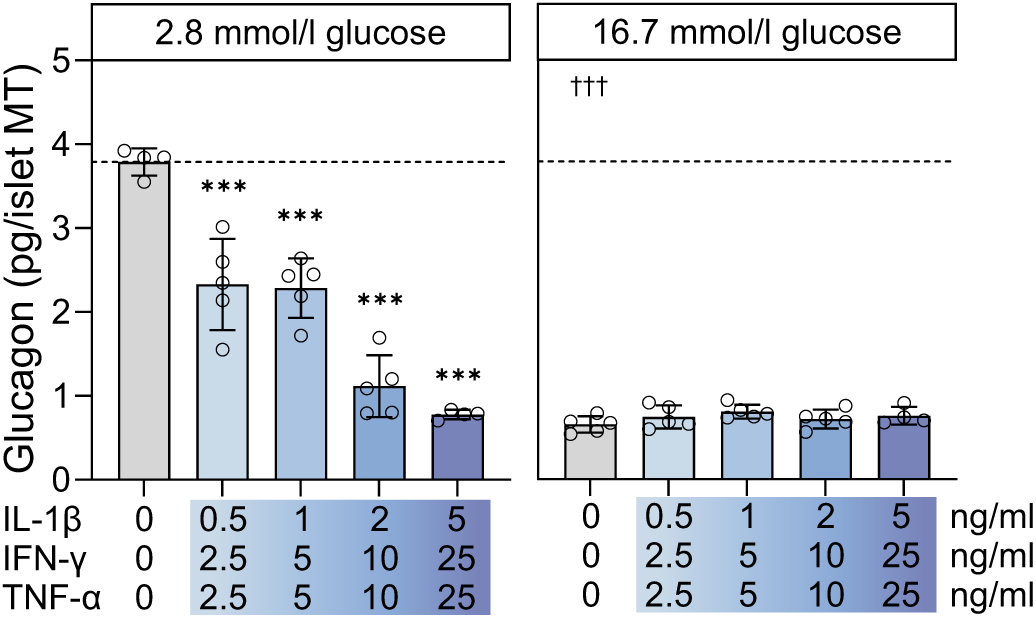
Related to Fig. 4. (a) Glucagon secretion at 2.8 and 16.7 mmol/l glucose in control (grey bars) and following long-term cytokine exposure with optimised dose titration (dose-¼, -½, -1 and -2). The dashed line denotes the untreated control baseline/physiological response to low glucose. Data presented as mean ± SD of donor 8 in 5 technical replicates. Asterisks (*) indicate statistical significance using one-way ANOVA with Dunnett’s multiple comparisons test compared to untreated control, ***p <0.001. Daggers (†) indicate statistical significance using Student’s t-test comparing the two untreated controls at 2.8 vs. 16.7 mmol/l glucose, †††p <0.001.

**ESM Fig. 5.**
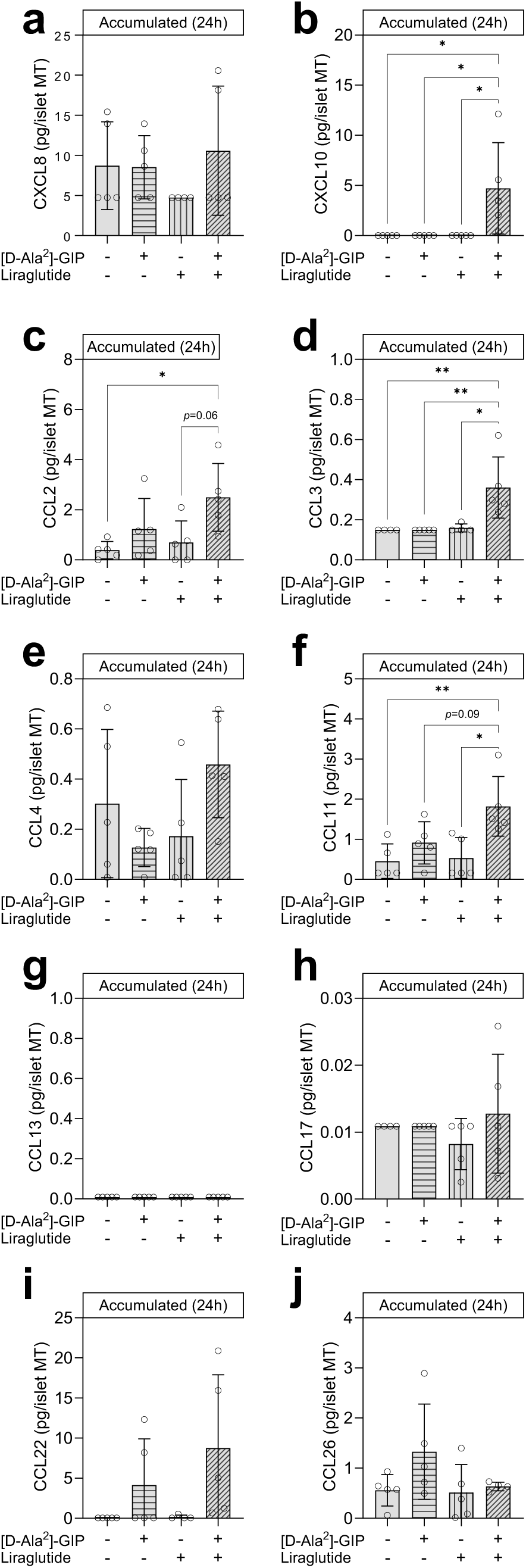
Related to Fig. 6. (a) Accumulated CXCL8, (b) CXCL10, (c) CCL2, (d) CCL3, (e) CCL4, (f) CCL11, (g) CCL13, (h) CCL17, (i) CCL22, and (j) CCL26 secretion during the last 24 hours of the untreated control (no bar pattern) or with 1 µmol/l [D-Ala^2^]-GIP (horizontal stripes), Liraglutide (vertical stripes), or [D-Ala^2^]-GIP+liraglutide (angled stripes). Data presented as mean ± SD of donor 11 in 5 technical replicates. Asterisks (*) indicate statistical significance using one-way ANOVA with Tukey’s multiple comparisons test, *p <0.05, **p <0.01.

